# Multi-echo BOLD fMRI improves cerebrovascular reactivity estimates in stroke

**DOI:** 10.64898/2026.02.03.703581

**Authors:** Rebecca G. Clements, Fatemeh Geranmayeh, Niamh V. Parkinson, Miguel Montero, Sebastian Urday, Katie Taran, Diego A. Caban-Rivera, Carson Ingo, Molly G. Bright

## Abstract

Cerebrovascular reactivity (CVR), the ability of cerebral blood vessels to dilate or constrict in response to a vasoactive stimulus, is a clinically meaningful measure of cerebrovascular health. Head motion and other noise sources substantially impact CVR quality, particularly in clinical populations. In this study, we evaluated multi-echo fMRI techniques, including optimal combination of echoes (ME-OC) and multi-echo independent component analysis (ME-ICA), for improving CVR quality relative to single-echo fMRI in participants with stroke. In a breath-hold fMRI dataset, ME-OC significantly improved CVR quality metrics and reduced the percentage of negative CVR values in normal-appearing gray and white matter (*p*<0.05). ME-ICA reduced the dependence of BOLD signals on head motion but did not improve CVR quality metrics. In a separate resting-state dataset, ME-OC effects were largely consistent with the breath-hold dataset, but ME-ICA also significantly improved CVR quality metrics and reduced negative CVR values in normal-appearing gray and white matter relative to ME-OC (*p*<0.05). These findings demonstrate that multi-echo fMRI can improve CVR estimation in clinical populations, particularly in low signal-to-noise datasets, enhancing the feasibility of CVR analyses in stroke studies and allowing for better visualization of stroke-related CVR deficits.

## Introduction

Cerebrovascular reactivity (CVR), the ability of cerebral blood vessels to dilate or constrict in response to a vasoactive stimulus, is an important measure of cerebrovascular health. CVR has clinical utility for a range of applications including brain tumor grading,^1^ the prediction of cognitive function,^2^ and the prediction of recurrent stroke risk.^3^ In participants with stroke, CVR could be used to guide personalized rehabilitation strategies^4^ and calibrate functional magnetic resonance imaging (fMRI) data to allow for more confident assessments of changes in neural activity.^5,6^ Several imaging approaches exist for assessing CVR,^7–9^ but blood-oxygen-level-dependent (BOLD) fMRI is most common. In a typical BOLD fMRI CVR experiment, arterial CO₂ is increased to trigger systemic vasodilation and increase cerebral blood flow. Arterial CO_2_ can be increased by intermittently inhaling air with a fixed concentration of CO_2_^10^. This approach has good inter-rater, inter-scanner, and test-retest reliability in healthy volunteers.^11^ Recent work has also shown that completing breath holds during fMRI can allow for robust mapping of CVR without the need for external gas delivery.^6,12–14^ CVR can even be quantified using resting-state fMRI by exploiting natural variations in arterial CO_2_ caused by fluctuations in breathing patterns.^9,15,16^

Despite the demonstrated success of these methods, BOLD CVR has not been widely adopted in clinical practice. A major methodological factor for this is the confounding effects of head motion in CVR estimation, limiting our ability to confidently assess meaningful changes in CVR. In fact, one study noted that head motion was the most common reason for a failed CVR examination.^17^ In studies using breath holds, an additional problem is task-correlated head motion, which further complicates the separation of motion-related signals from signals of interest.^18^ With resting-state fMRI approaches, head motion can diminish the detectability of BOLD fluctuations related to spontaneous fluctuations in arterial CO_2_, which make up no more than 20% of the overall signal variance.^16^

Several denoising methods have been proposed to mitigate the effects of head motion on fMRI. Most commonly, each volume in the timeseries is realigned to a reference volume. To further compensate for motion-related signal changes in the data, the timeseries of the 6 estimated motion parameters and their derivatives can be used as nuisance regressors in a regression model.^19^ Censoring can also be applied to remove time points with high levels of head motion;^20,21^ however, censoring reduces the temporal degrees of freedom and cannot fully account for prolonged or step-like signal changes related to motion.^22^ Data-driven techniques such as principal component analysis and independent component analysis can also be used to identify and remove components that are related to motion;^23–25^ however, many of these methods cannot fully remove head motion artifacts,^26^ especially when signals of interest are correlated with motion.^27^

An alternative approach to reduce noise in fMRI involves a multi-echo acquisition, in which multiple echo times are acquired for each MRI volume. This approach enables voxel-wise estimation of T_2_*, allowing echoes to be combined by weighted averaging to maximize sensitivity and reduce random noise, particularly in brain regions prone to strong susceptibility artifacts.^28,29^ Motion-related artifacts can further be mitigated by applying multi-echo independent component analysis (ME-ICA). ME-ICA involves decomposing the data into independent components and then classifying them as BOLD signals, reflecting changes in the transverse relaxation rate (R_2_*), or non-BOLD signals, reflecting changes in initial signal intensity (S_0_). The latter arises from factors such as subject motion, venous sinus blood flow, cardiac pulsation, and field fluctuations.^30^ This classification is based on the kappa and rho parameters for each component, which reflect the extent to which signal changes agree with ΔR_2_* and ΔS_0_ signal models.^30^ By leveraging echo-time dependence, ME-ICA enables better identification of noise components compared to conventional, single-echo ICA approaches.^31^

Compared to single-echo fMRI, optimal combination of echoes has been shown to be superior for detecting neuronal activation^32,33^ and to improve repeatability, sensitivity, specificity, and reliability of breath-hold BOLD CVR estimates in healthy participants.^34^ For mapping motor activation in healthy participants, ME-ICA has shown additional benefits over using optimal combination alone, allowing for better disassociation of the effects of head motion on the BOLD signal.^32^ ME-ICA has also demonstrated benefits for precision functional mapping of intrinsic connectivity in resting-state fMRI, as it improves test-retest reliability and reduces the need for multiple long scans.^35^ Most relevant, a recent study examined how ME-ICA strategies affect CVR estimates in healthy participants and found that a “conservative” ME-ICA approach yielded the largest reduction of motion-related artifacts while producing reliable CVR estimates.^18^

As previous studies evaluating the benefits of multi-echo fMRI for CVR estimates have been limited to healthy participants, the goal of the current study is to evaluate the benefits of multi-echo fMRI techniques for estimating CVR in participants with stroke. Since participants with stroke tend to exhibit higher levels of head motion compared to controls,^36^ ME-ICA may be particularly advantageous for uncoupling BOLD signal fluctuations from motion confounds and improving the quality of subsequent CVR maps. Additionally, by reducing false positive and false negative CVR estimates, ME-ICA may aid in the delineation of brain regions with true, stroke-related CVR deficits. In this study, we aim to evaluate how optimal combination of echoes (ME-OC), with and without ME-ICA, affects CVR quality and sensitivity to stroke-related pathology compared to the conventional single echo approach. We evaluate these methods using a breath-hold fMRI dataset consisting of 64 participants with stroke. We then assess the generalizability of our findings to a resting-state fMRI dataset for a separate cohort of 27 participants with stroke.

## Methods

### Breath-hold data collection

64 participants with stroke (49 M, 61 ± 10 years) were imaged at Imperial College London on a 3T Siemens Magneton Verio scanner 137 ± 71 (mean ± SD) days after stroke onset. The data were acquired as part of a longitudinal observational clinical study approved by UK’s Health Research Authority (Registered under NCT05885295; IRAS:299333).^37^ This study was carried out in strict compliance with national legislation and the General Data Protection Regulation and all participants gave written, informed consent. Of the 64 participants, 52 had ischemic strokes and 12 had hemorrhagic strokes. Inclusion and exclusion criteria for this study have been previously described by Gruia and colleagues.^37^ A high-resolution 1mm isotropic T1-weighted whole brain structural image was acquired, followed by the acquisition of a 1mm isotropic fluid-attenuated inversion recovery (FLAIR) image. BOLD fMRI data were acquired using a multi-band multi-echo gradient-echo echo planar imaging sequence (TR = 1.792s, TEs = 13.6/26.97/40.34/53.71ms, voxel size = 3mm isotropic, MB factor = 2, 154 volumes). During the scan, participants completed a breath-hold task^6^ that consisted of 6 repetitions of the following paradigm: 14 seconds of natural breathing, 16 seconds of paced breathing, and a 15 second end-expiration breath hold. Expired CO_2_ was measured during the scan at a sampling frequency of 1000 Hz using a nasal cannula attached to a gas analyzer. In addition, a gradient-echo field map was acquired to be used for distortion correction (TR = 622ms, TEs = 4.92/7.38ms, FA = 50°, voxel size = 2.39 × 2.39 × 3.00 mm^3^).

### Breath-hold data analysis

Expired CO_2_ data were processed using an in-house MATLAB script. After a peak-detection algorithm identified end-tidal peaks, the results of the algorithm were manually verified, and the peaks were linearly interpolated to create P_ET_CO_2_ timeseries with the same frequency as the original CO_2_ data. Next, P_ET_CO_2_ timeseries were convolved with the canonical hemodynamic response function^38^ and converted to units of mmHg. As high-quality P_ET_CO_2_ timeseries are required for reliable CVR calculation,^12,39,40^ we evaluated the quality of the P_ET_CO_2_ timeseries by computing the relative power in the dominant frequency range of the breath-hold task (0.0192 to 0.0252 Hz).^39^ P_ET_CO_2_ timeseries with less than 50% power in the BH range were classified as low-quality and excluded them from further analyses.^39^

Masks of stroke lesions and white matter hyperintensities, which we will refer to collectively as vascular lesions, were manually delineated using the FLAIR images and verified by neurologist FG. Pre-processing of the T1-weighted and fMRI images was performed using custom scripts available at https://github.com/BrightLab-ANVIL/PreProc_BRAIN which utilized both FSL^41^ and AFNI^42,43^ tools. The T1-weighted images were brain-extracted with *BET*^41^ and then segmented into gray matter, white, matter and cerebrospinal fluid (CSF) with *FAST*^44^ using a threshold of 0.5. Vascular lesions were transformed to T1-space using *FLIRT*^45,46^ and then subtracted from the gray and white matter masks to generate masks of normal-appearing gray and white matter. For each echo of the functional data, head-motion realignment was performed with AFNI’s *3dVolReg* using the 1^st^ volume of the 1^st^ echo as the reference volume and then distortion correction was performed with *FUGUE*.^47,48^ A brain mask was generated using the distortion-corrected reference volume with *BET*^45^ and used to brain-extract each echo.

### Generation of single-echo (SE) datasets

The pre-processed second echo data (TE = 26.97ms) was used as a surrogate for single-echo data. Spatial smoothing was applied with a 4mm full width at half maximum (FWHM) Gaussian kernel using AFNI’s *3dmerge*.

### Generation of multi-echo optimally combined (ME-OC) datasets

*Tedana*^29,30,49,50^ was used to generate a weighted average of the 4 pre-processed echoes based on T_2_* estimates, producing multi-echo optimally combined (ME-OC) datasets. Spatial smoothing was applied with a 4mm FWHM kernel.

### Multi-echo independent component analysis (ME-ICA)

When running *tedana* to generate the ME-OC datasets, multi-echo independent component analysis (ME-ICA) was performed using the demo_external_regressors_single_model decision tree included in *tedana* v24.0.2 and described in the tedana documentation.^51^ This decision tree uses information from the kappa and rho parameters to determine if components are likely to be BOLD (signal) or non-BOLD (noise). It also classifies components as noise if they significantly fit a model of external nuisance regressor time series and the fit models a substantial amount of the total variance. Nuisance regressors used were motion parameters from volume registration and their derivatives. A similar component classification strategy has been previously implemented by Reddy and colleagues.^52^

### Breath-hold CVR calculation

3 CVR maps (SE, ME-OC, and ME-ICA) were generated for each subject using *phys2cvr*^53^ with a lag range of ±12 seconds in 0.3 second increments. *phys2cvr* is a lagged general linear modeling (GLM) approach that allows for simultaneous modeling of nuisance and (shifted) P_ET_CO_2_ regressors to account for voxel-wise hemodynamic delays and calculate CVR.^54^ The SE and ME-OC data were modeled using a design matrix consisting of the shifted P_ET_CO_2_ timeseries convolved with the HRF, Legendre polynomials up to 3^rd^ order, and the 6 demeaned motion parameters with each of their temporal derivatives. For the ME-ICA maps, following the conservative ME-ICA approach described previously,^18^ we additionally modeled the ME-OC data using ICA noise components orthogonalized to the shifted P_ET_CO_2_ timeseries convolved with the HRF, ICA signal components, motion parameters and their derivatives, and Legendre polynomials. These methods are summarized in Figure 1. For visualization purposes only (the calculation of all quality metrics described below was performed in native space), we registered each whole-brain CVR amplitude map to MNI space using the FSL 1mm MNI template resampled to 3 mm resolution (*FLIRT* and *FNIRT*, FSL).

**Figure 1.**
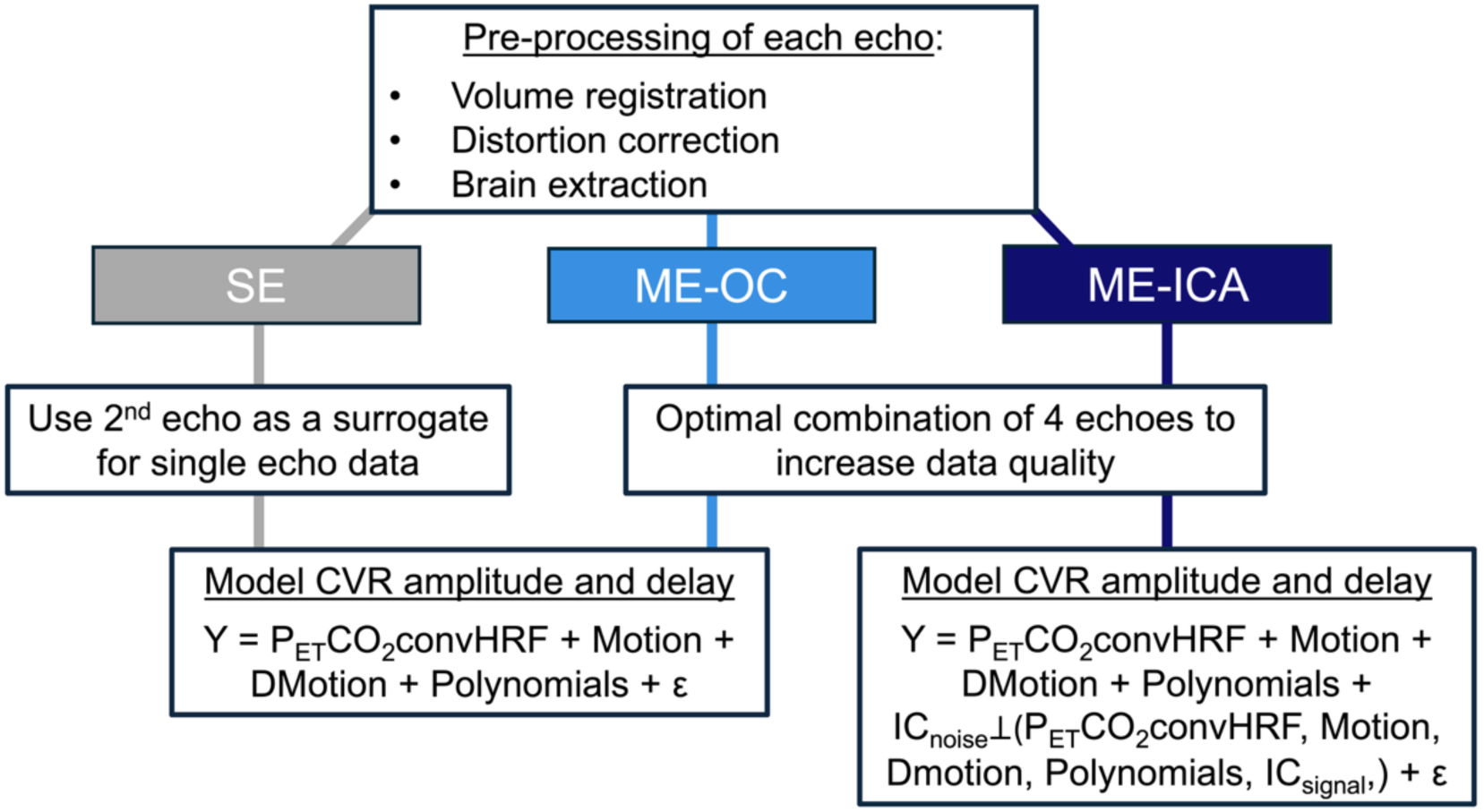
Overview of the methods for generating breath-hold CVR maps using the SE, ME-OC, and ME-ICA methods. The SE approach uses the smoothed, pre-processed 2^nd^ echo as a surrogate for single-echo data, while the ME-OC and ME-ICA approaches use the smoothed optimal combination of the 4 pre-processed echoes to increase data quality. The SE and ME-OC approaches model CVR using the shifted end-tidal CO_2_ regressor convolved with the HRF (P_ET_CO_2_convHRF), motion parameters, motion derivatives (DMotion), and Legendre polynomials up to 3^rd^ order. The ME-ICA approach additionally models CVR using the ME-ICA components classified as noise orthogonalized to P_ET_CO_2_convHRF, motion parameters, DMotion, Legendre polynomials, and the ME-ICA components classified as signal.

### Assessment of the impact of head motion on BOLD signals

To quantify head motion during the breath-hold fMRI scan, framewise displacement (FD) was calculated as the sum of the absolute difference of the volume realignment parameters (see Reddy and colleagues^32^ for details).^55^ DVARS was calculated as the spatial root mean square of the first derivative of the denoised signal.^56^ Denoised datasets were obtained for each method by subtracting the fitted timeseries associated with the nuisance regressors (i.e., all regressors shown in Figure 1 other than P_ET_CO_2_ convolved with the HRF for each method) from the unsmoothed single-echo or optimally combined fMRI dataset (*3dREMLfit*, AFNI). For each method, DVARS was calculated (*3dTto1D*, AFNI) within the tighter brain mask created after running *tedana*.^32^

Next, the slope of DVARS vs. FD (DVARS ∼ FD) was calculated for each scan and method. We hypothesized that the datasets denoised using ME-ICA would have a lower association between head motion and BOLD signals than the ME-OC and SE datasets. To test this hypothesis and to verify that ME-OC did not affect the relationship between head motion and BOLD signals compared to SE, we conducted one-sided paired *t*-tests. These tests evaluated whether the slope significantly decreased when using ME-OC compared to SE, ME-ICA compared to SE, and ME-ICA compared to ME-OC (significance threshold *p*<0.05, with Bonferroni correction).

### Breath-hold CVR quality metrics

To assess whether ME-OC and ME-ICA increased the detectability of true CVR responses relative to SE, we evaluated if they increased the percentage of significant voxels in normal-appearing gray and white matter. CVR values were considered significant if they had absolute T-statistics greater than 1.98 (corresponding to *p*<0.05). One-sided paired *t*-tests were conducted to assess significant increases when using ME-OC compared to SE, ME-ICA compared to SE, and ME-ICA compared to ME-OC (significance threshold *p*<0.05, with Bonferroni correction).

Additionally, we wanted to evaluate whether ME-OC and ME-ICA significantly increased spatial coherence in normal-appearing gray and white matter. We expected more effective denoising to make neighboring voxels of the same tissue type to have more similar CVR values. Moran’s *I* is a measure of spatial coherence that ranges from -1 to +1; more positive values indicate that neighbors have similar values, while more negative values indicate that neighbors have dissimilar values.^57^ To calculate Moran’s *I*, we created a nearest neighbor weights matrix based on 26 nearest neighbors using the libpysal Python library and then used the Moran function in the ESDA Python library.^58^ 1-sided paired *t*-tests were conducted to evaluate whether Moran’s *I* significantly increased when using ME-OC compared to SE, ME-ICA compared to SE, and ME-ICA compared to ME-OC (significance threshold *p*<0.05, with Bonferroni correction).

### Resting-state data collection

To assess the generalizability of our findings to resting-state CVR estimates, we analyzed a dataset consisting of 27 participants with ischemic stroke (19 M, 65±12 years), scanned approximately 4 weeks after stroke onset on a 3T Siemens Prisma MRI scanner at Northwestern University. This study was approved by the Northwestern University Institutional Review Board and was conducted in accordance with the ethical guidelines in the Belmont Report. All participants provided written, informed consent. After acquiring a T1-weighted whole-brain MPRAGE and a FLAIR, fMRI data were acquired using a multi-band, multi-echo gradient-echo echo planar imaging sequence (TR = 1.5s, TEs = 10.6/27.84/45.06/62.3/79.52ms, voxel size = 2.5mm isotropic, 287 volumes). During the fMRI scan, participants were instructed to relax and focus on a fixation cross. For distortion correction and to facilitate functional realignment, one single-band reference volume (SBRef) was also acquired before the functional acquisition using the same scan acquisition parameters without the multi-band acceleration factor. P_ET_CO_2_ recordings were not acquired due to the need to keep the scan setup time as short as possible.

### Resting-state fMRI data processing

Masks of acute infarcts, hereafter referred to as vascular lesions, were delineated using acute diffusion-weighted MRI scans. fMRI pre-processing steps were as similar as possible to those described for the breath-hold dataset (see Supplementary Material for details on infarct identification and fMRI pre-processing). As we did not acquire a P_ET_CO_2_ regressor during the scan and could not map CVR using the same methods as the breath-hold data, we chose to use the *Rapidtide* toolbox^59^ to compute a surrogate measure related to CVR. *Rapidtide* performs cross correlations between a reference timeseries and every voxel in an fMRI dataset to produce an R^2^ map. The reference timeseries should reflect systemic physiological fluctuations;^60^ to avoid brain areas that may have abnormal perfusion dynamics,^39^ we used the mean time series in gray matter in the non-lesion hemisphere as the reference. One subject had bilateral lesions, and we used the less affected hemisphere as the reference for this subject. The reference timeseries is iteratively refined to better reflect the hemodynamic component of the BOLD signal, and the R^2^ map reflects the degree to which the blood signal corresponds to the BOLD signal in each voxel.^61^ As described by Donahue and colleagues,^61^ conditions with altered CVR will have decreased R^2^ in the affected voxels compared to controls, and thus voxel-wise R^2^ values are a reasonable surrogate for CVR. Because the R^2^ map does not indicate the direction of the relationship between the reference timeseries and each voxel’s time series, we also provide maps of the maximum absolute correlation, which preserve the sign of the correlation, to be interpreted alongside the R^2^ maps.

Prior to running *Rapidtide*, we obtained smoothed, denoised datasets (SE, ME-OC, and ME-ICA) for each method using similar design matrices as those in Figure 1 except we could not include P_ET_CO_2_ regressors (see Supplementary Material for details). Then, for each method, we ran *Rapidtide* to model each dataset using a shift range of -12 to 12 seconds and the –bipolar option to allow for both positive and negative correlation values.

FD was calculated using the methods described for the breath-hold dataset, and then DVARS was calculated from the denoised, unsmoothed datasets (*3dTto1D*, AFNI). To better understand how ME-OC and ME-ICA affect the fit of each voxel’s timeseries with that of the probe regressor, we calculated the average R^2^ in gray and white matter. We also assessed spatial coherence by calculating Moran’s *I* in normal-appearing gray and white matter, using the same methods described for the breath-hold dataset. For the visualization of 3 representative subjects in this manuscript, we registered their maps to MNI space and generated masks of white matter injuries; see Supplementary Material for details.

## Results

### Evaluation of P_ET_CO_2_ timeseries in the breath-hold dataset

28 participants were excluded due to having less than 50% power in the dominant frequency range of the breath-hold task. The resulting breath-hold dataset size was 36 participants (28 M, 10 participants with hemorrhagic stroke, 63 ± 12 years). See Supplementary Material for a justification of the sample size. It is important to note that while this level of attrition is conservatively high, the goal of this study was not to evaluate the feasibility of breath-hold tasks but to assess whether multi-echo approaches improve breath-hold CVR estimates in stroke.

### Assessment of the impact of head motion on BOLD signals in the breath-hold dataset

The relationship between FD and DVARS for one example subject in the breath-hold dataset is illustrated in Figure 2A. While the best-fit lines for the SE and ME-OC approach appear relatively similar, the ME-ICA approach exhibits a weaker association between DVARS and FD, suggesting that ME-ICA decreases the impact of head motion on BOLD signals. This relationship is consistent across all of the subjects: the ME-ICA approach resulted in significantly decreased slopes of DVARS versus FD compared to the SE and ME-OC approaches (Figure 2B; *p*<0.05, Bonferroni corrected). As expected, we did not observe a significant decrease in the slope of DVARS versus FD when using ME-OC compared to SE.

**Figure 2.**
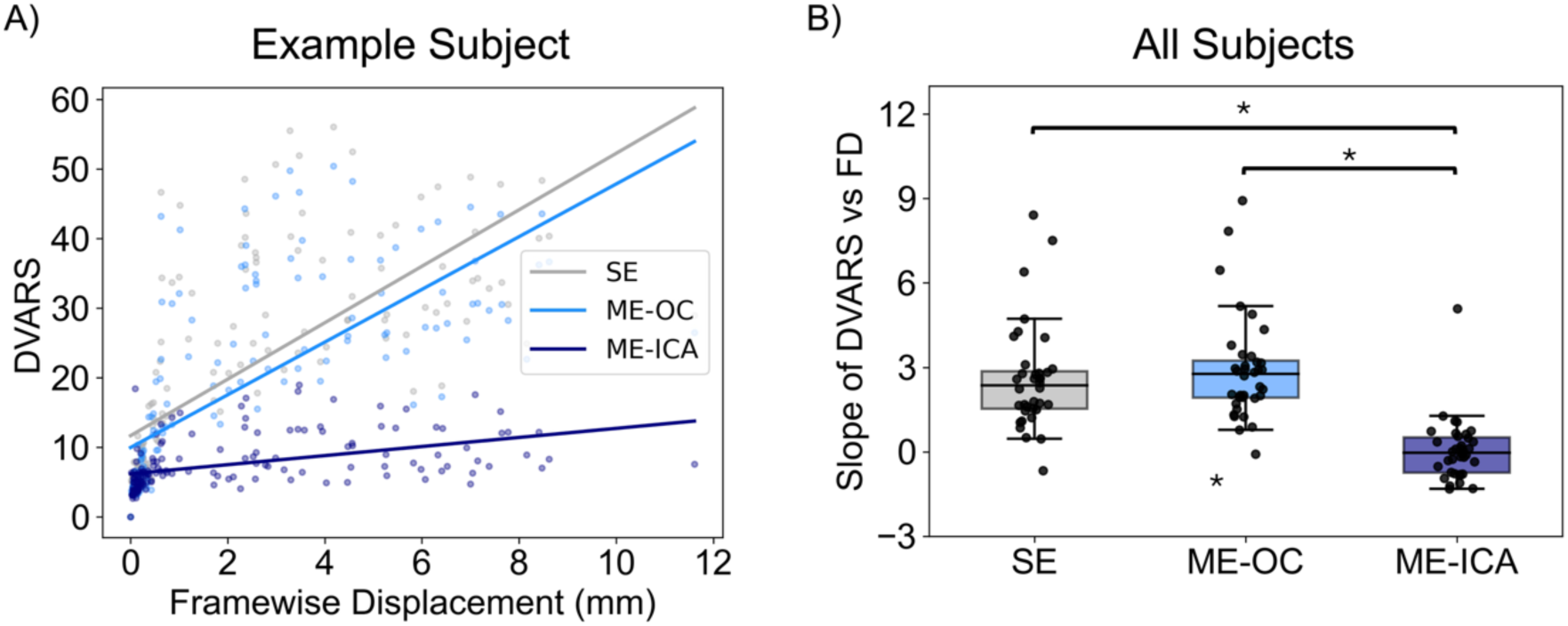
DVARS versus framewise displacement for a single example participant (left) and the slopes of DVARS versus FD across all participants (right). The slope of DVARS versus FD represents the extent to which head motion impacts BOLD signals. In the left plot, each point represents a timepoint and each line represents the linear regression for the entire timeseries. Asterisks indicate significant differences (*p*<0.05, Bonferroni corrected).

### Breath-hold CVR quality metrics

Compared to SE, ME-OC and ME-ICA significantly increased the percentage of normal-appearing gray and white matter voxels with significant CVR (Figure 3A; *p*<0.05, Bonferroni corrected). We did not observe a significant increase in the percentage of voxels with significant CVR with using ME-ICA compared to ME-OC. Similarly, ME-OC and ME-ICA significantly increased spatial coherence in gray and white matter compared to SE (*p*<0.05, Bonferroni corrected), but ME-ICA did not significantly increase spatial coherence compared to ME-OC (Figure 3B).

**Figure 3.**
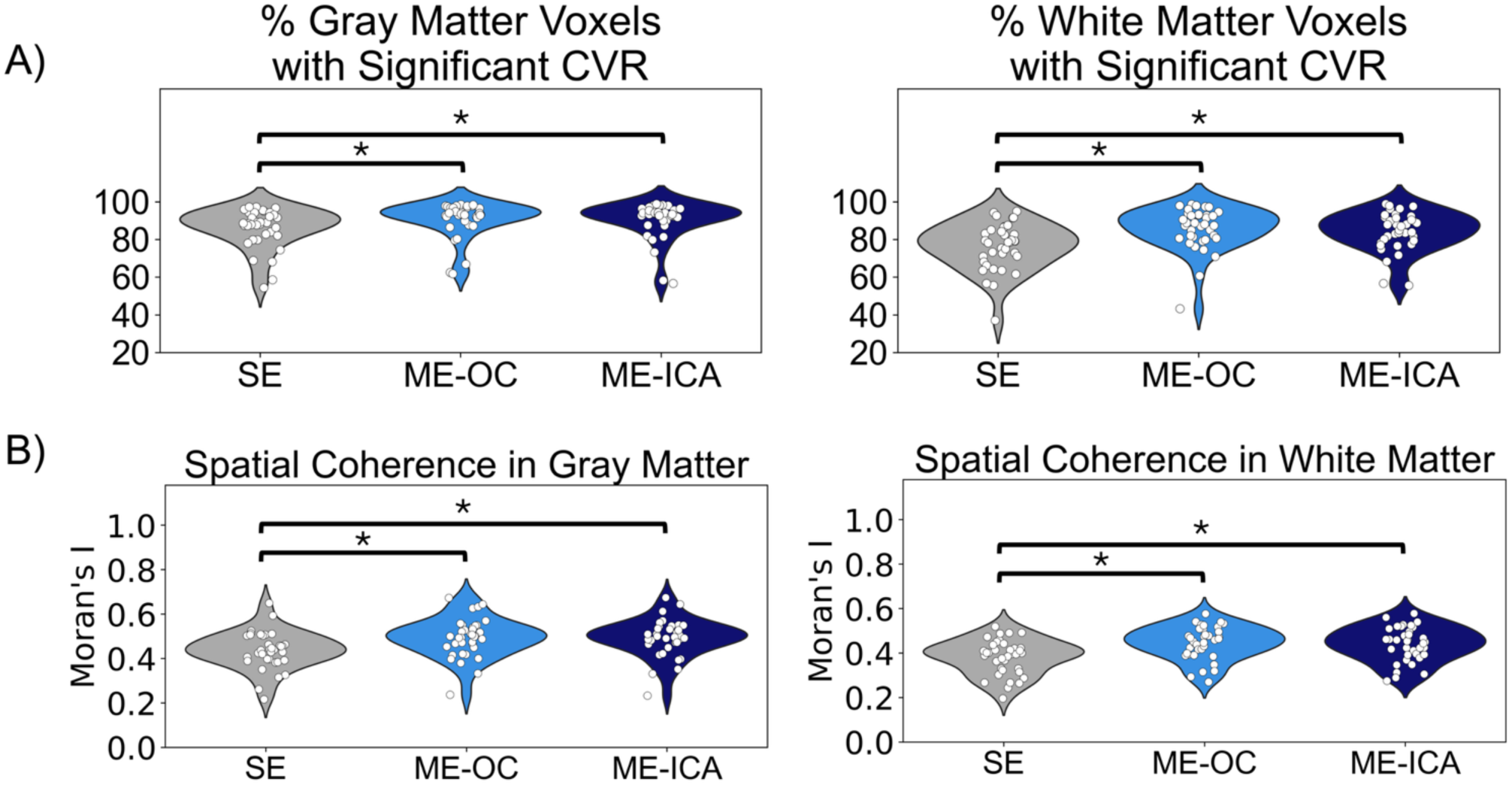
Plots showing changes in the percentage of normal-appearing gray and white matter voxels with significant CVR (A) and spatial coherence in normal-appearing gray and white matter (B) across the 3 methods. Asterisks indicate significant differences (*p*<0.05, Bonferroni corrected).

### Example breath-hold CVR amplitude maps

Compared to the SE approach, the ME-OC approach produces CVR amplitude maps with negative values that are more confined to vascular lesions, increasing confidence that these negative values reflect meaningful physiological phenomena (Figure 4). For instance, in Example 1 of Figure 4, the SE map shows a cluster of negative CVR values in the anterior white matter, distant from lesions (light green arrow). Many of these voxels become positive in the ME-OC map, while the negative voxels located within or adjacent to lesions remain unchanged. Similarly, in Example 2, the SE map contains widespread negative values throughout the anterior portion of the brain, making it difficult to distinguish stroke-related CVR abnormalities from artifacts related to noise. In the ME-OC map, many of these spurious negative voxels disappear, allowing the negative values surrounding the lesion to be more clearly delineated. Across all of the subjects, this effect was less pronounced when comparing ME-ICA to ME-OC, but the ME-ICA approach did make negative CVR values more confined to vascular lesions in some subjects (see Examples 4 and 6 in Figure 4).

**Figure 4.**
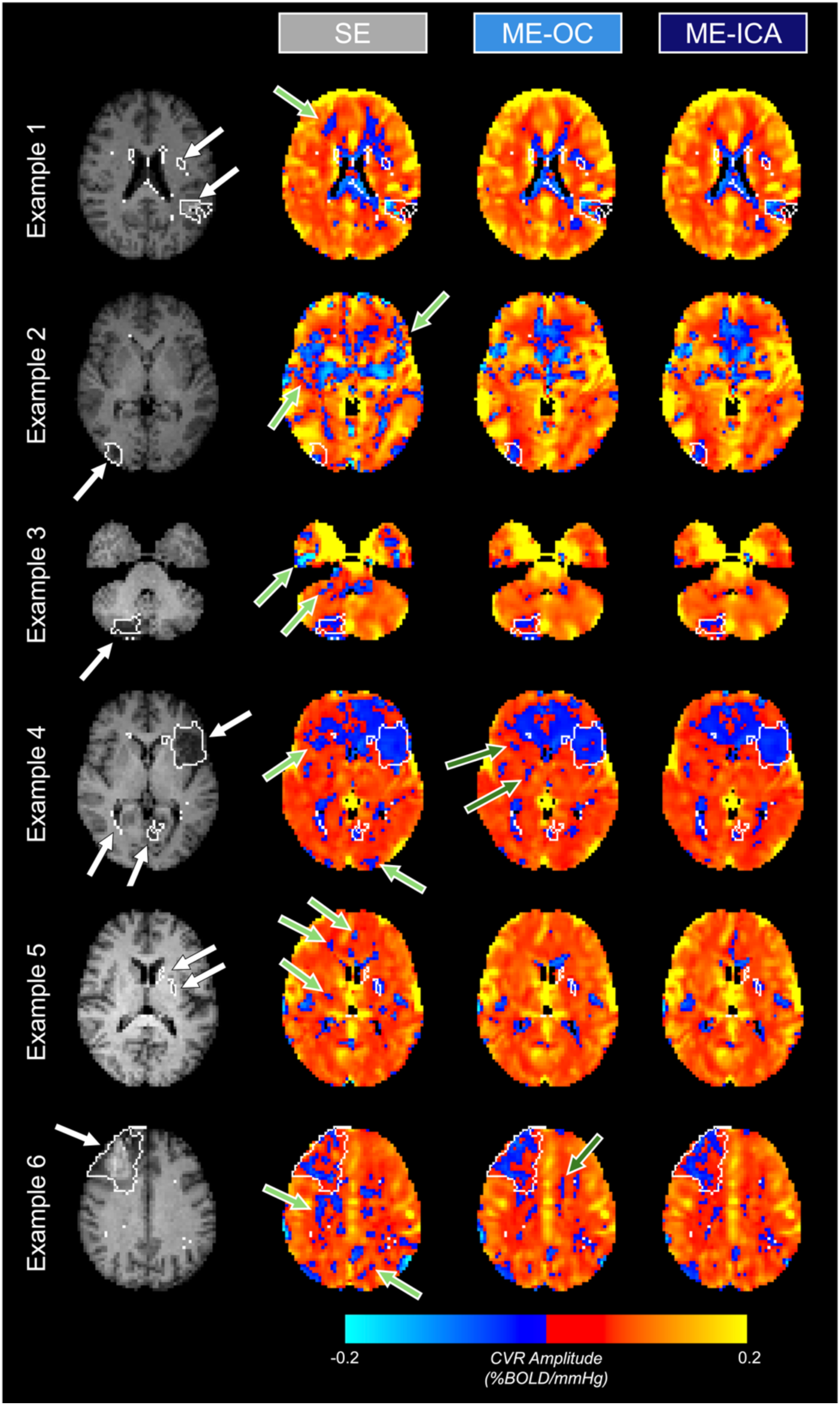
SE, ME-OC, and ME-ICA CVR amplitude maps for 6 example subjects. For reference, T1-weighted images for each example subject are shown in the first column. Compared to SE, ME-OC makes negative CVR values more confined to vascular lesions, outlined in white and indicated by white arrows (first column). Light green arrows (second column) indicate non-lesion areas with fewer negative CVR values using ME-OC compared to SE. Dark green arrows (third column) indicate non-lesion areas with fewer negative CVR values using ME-ICA compared to ME-OC.

### Assessment of the generalizability of findings to resting-state CVR estimates

In the resting-state dataset, we observed the same changes in the slope of DVARS versus FD that we observed in the breath-hold dataset: the ME-ICA approach resulted in a significant decrease of the slope compared to the SE and ME-OC approaches (*p*<0.05, Bonferroni corrected), indicating that ME-ICA effectively removes motion-related signal changes (Figure 5A). In both gray and white matter, the ME-OC approach significantly increased R^2^ compared to SE, and ME-ICA resulted in an additionally significant increase in R^2^ compared to ME-OC (Figure 5B; *p*<0.05, Bonferroni corrected). This effect, reflecting progressive improvements in model fits with ME-OC and ME-ICA, is also evident in the example maps shown in Figure 6. In gray matter, the ME-OC approach significantly increased spatial coherence compared to SE (*p*<0.05, Bonferroni corrected), but the ME-ICA approach did not significantly increase spatial coherence compared to ME-OC (Figure 5C). However, in white matter, not only did the ME-OC approach significantly increase spatial coherence compared to SE, but the ME-ICA approach also significantly increased spatial coherence compared to ME-OC (*p*<0.05, Bonferroni corrected). We again observed that ME-OC and ME-ICA make negative correlation values more confined to vascular lesion areas (Figure 6).

**Figure 5.**
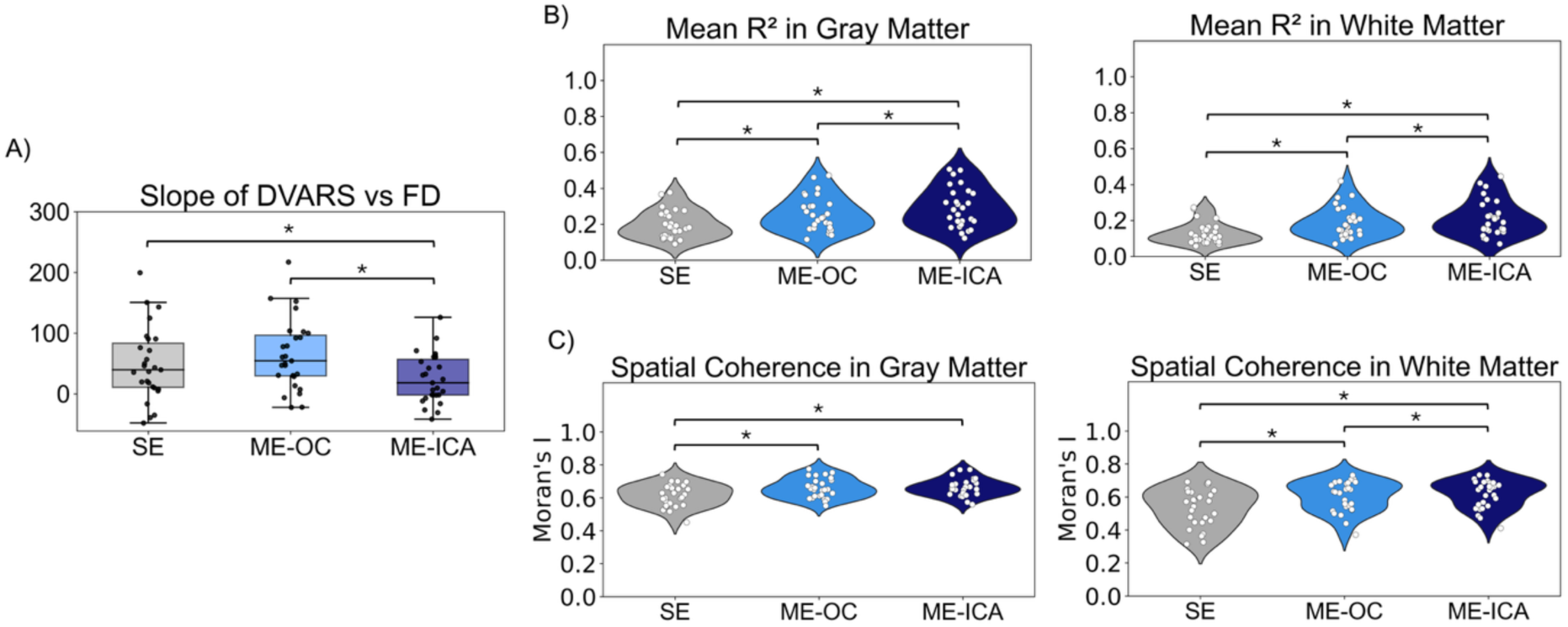
Slope of DVARS versus FD (A), mean R^2^ in normal-appearing gray and white matter (B), and spatial coherence in normal-appearing gray and white matter (C) for resting-state datasets. Asterisks indicate significant differences (*p*<0.05, Bonferroni corrected).

**Figure 6.**
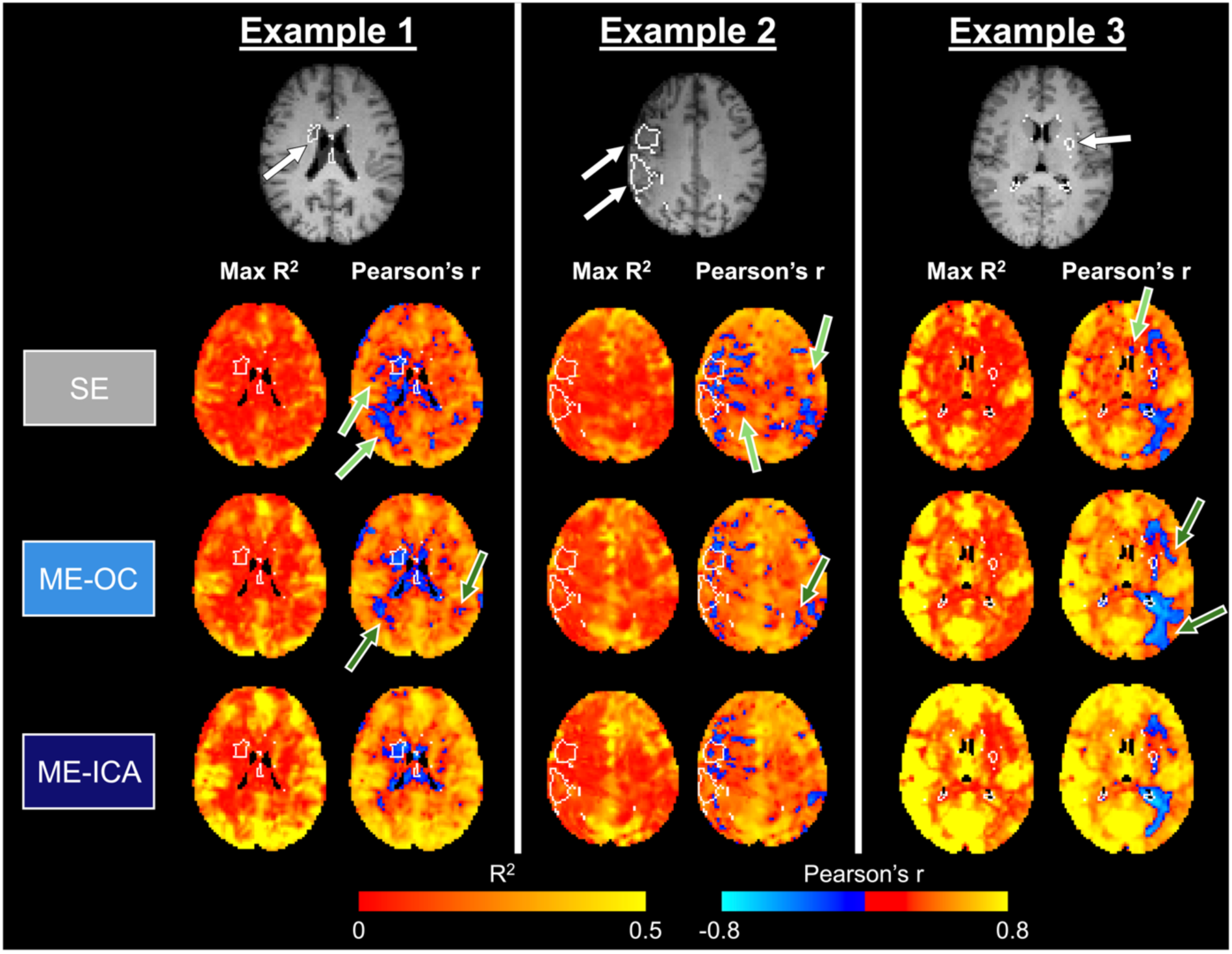
Maps of the maximum R^2^ and r for 3 example resting-state datasets. R^2^ acts as a surrogate measure for CVR, while r provides complementary information about the direction of the correlation. Vascular lesions are outlined in white and indicated by white arrows on the T1-weighted anatomical scan (first row). Light green arrows (second row) indicate non-lesion areas with fewer negative correlation values using ME-OC compared to SE. Dark green arrows (third row) indicate non-lesion areas with fewer negative correlation values using ME-ICA compared to ME-OC.

## Discussion

In this study, we demonstrated the utility of multi-echo fMRI for improving CVR estimates in participants with stroke. This work complements ongoing efforts to improve the feasibility of breath-hold fMRI for CVR mapping in clinical populations. For example, recent work aimed at improving breath-hold feasibility demonstrated that 9-second breath-hold periods, shorter and more tolerable than the commonly used 15-second breath holds, yield highly reproducible CVR estimates.^62^ Prior studies have also explored alternative approaches for mapping CVR when P_ET_CO_2_ measurements are unreliable due to challenges with breath-hold task compliance, including methods that leverage information from a respiration belt.^39,40^ Because respiration belt recordings were not available in the present study, over 40% of participants were excluded due to low-quality P_ET_CO_2_ timeseries. This level of attrition underscores the need for a combination of advanced translational methods to mitigate challenges such as task compliance and head motion to make CVR estimation in clinical populations more feasible.

Building on these efforts, we evaluated multi-echo fMRI methods for mitigating the effects of head motion and other noise sources on CVR quality in participants with stroke. The incorporation of multi-echo fMRI into clinical imaging pipelines is increasingly feasible: methodological advances such as multiband acquisition enable minimal loss of spatial and temporal resolution relative to single-echo fMRI,^63^ and publicly available software packages provide automated processing of multi-echo data.^42,51^ We showed that ME-OC resulted in clear quantitative improvements to CVR quality in both breath-hold and resting-state data. While further adding ME-ICA significantly reduced motion-related signal fluctuations in both datasets, it only yielded quantitative CVR quality improvements in resting-state data, suggesting that ME-ICA may be most valuable in low signal-to-noise datasets. Together, these results, discussed in more detail below, motivate the implementation of multi-echo fMRI strategies in future clinical CVR studies.

### While ME-OC shows consistent benefits across datasets, ME-ICA may be most beneficial in lower-signal and higher-motion datasets

In the SE CVR amplitude maps, many subjects exhibited clusters of negative voxels in normal-appearing gray and white matter that became positive with the ME-OC approach, suggesting that these negative values may have been noise-related (Figures 4 and 6). To quantify this effect at the group level, we performed a post hoc analysis to compute the percentage of normal-appearing gray and white matter voxels with negative CVR for each method (Figure 7). Consistent with our qualitative observations, ME-OC significantly decreased the percentage of normal-appearing gray and white matter voxels with negative CVR values relative to SE in both datasets. In the resting-state dataset, ME-ICA further reduced the percentage of normal-appearing gray and white matter voxels with negative CVR values relative to ME-OC, demonstrating that ME-ICA may be particularly valuable in datasets with low signal-to-noise ratios. Together, these findings indicate that multi-echo fMRI techniques result in more plausible CVR values in normal-appearing tissue and may aid in distinguishing noise effects from physiologically meaningful CVR alterations.

**Figure 7.**
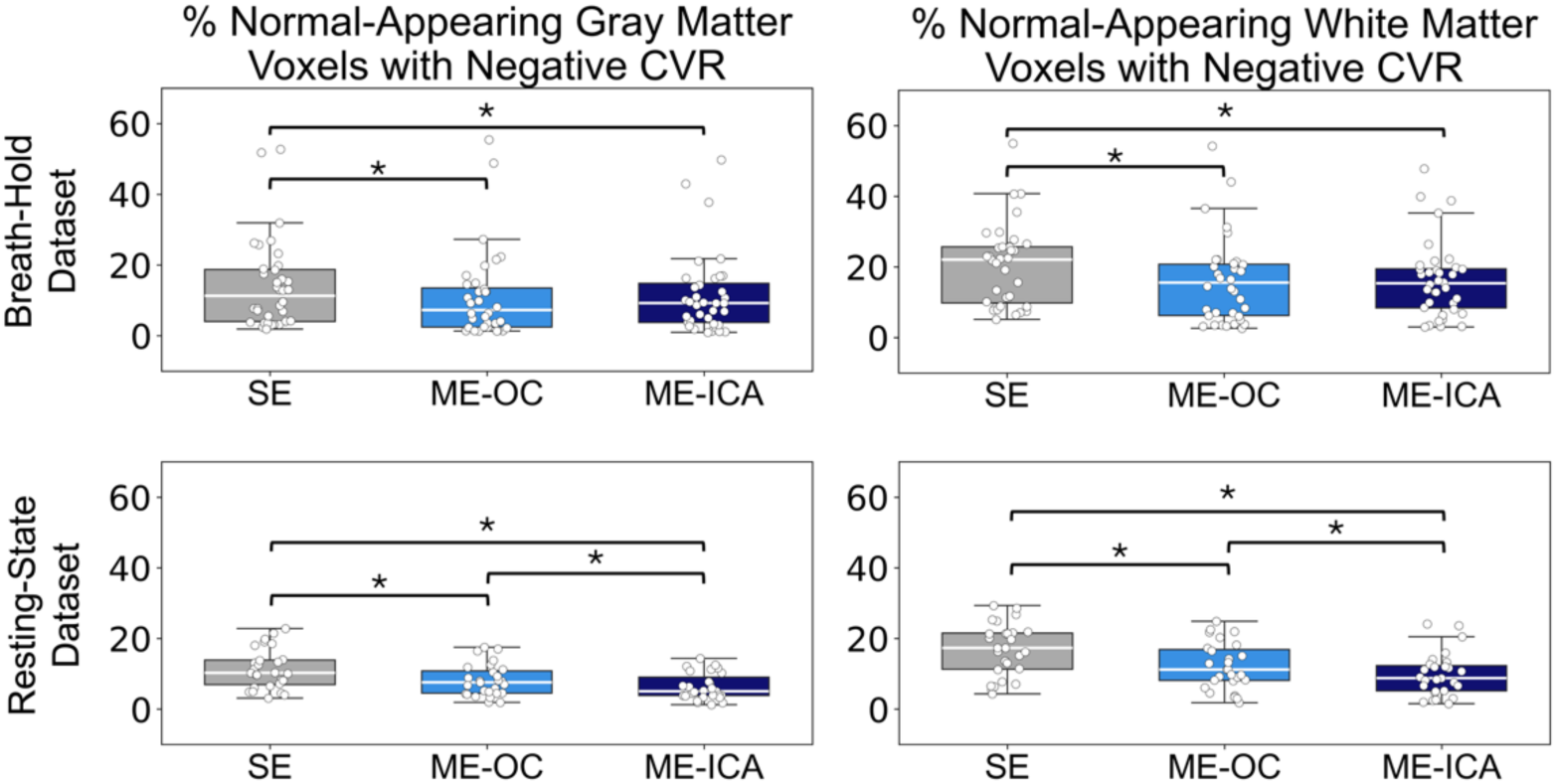
Percentage of normal-appearing gray and white voxels with negative CVR values in the breath-hold and resting-state datasets. Note that for this analysis in the resting-state dataset, we used the maps of Pearson’s *r* as a surrogate for CVR. Asterisks indicate significant differences determined using a 2-sided paired *t*-test (significance threshold *p*<0.05, with Bonferroni correction).

We also evaluated the effects of ME-OC and ME-ICA on CVR quality metrics relative to SE. ME-OC consistently improved CVR quality, evidenced by an increased percentage of voxels with significant CVR in the breath-hold dataset and higher mean R^2^ in the resting-state dataset, as well as increased spatial coherence in both datasets (Figures 4 and 6). On the other hand, while ME-ICA made BOLD signals less dependent on head motion in both the breath-hold and resting-state datasets (Figure 2), it only resulted in significant CVR quality improvements in the resting-state dataset. Again, this is likely related to the lower signal-to-noise-ratio of vascular responses in the resting-state data compared to the breath-hold data, making resting-state CVR estimates more vulnerable to motion effects and other non-BOLD artifacts.^64,65^

In this context, we would also expect ME-ICA to be most beneficial in participants with higher motion, and thus higher noise levels. To better understand whether this was the case, we qualitatively examined CVR maps generated using SE, ME-OC, and ME-ICA for the 3 participants with the highest motion levels during scanning (Figure S1). The participant with the highest average FD (2.18mm) exhibited more than 3.5 times more motion than the participant with the second-highest average FD, and the CVR maps appeared highly noisy and similar across the 3 methods. This suggests that when motion levels are extremely high, high-quality CVR maps may not be salvageable, even with advanced denoising methods. For the 2 subjects with the second highest motion levels (FD = 0.62 and 0.61), ME-ICA showed clear benefits, removing clusters of negative CVR outside of vascular lesion areas. Together, these findings indicate that ME-ICA can meaningfully improve CVR estimates in high-motion data, but its benefits are contingent on data quality remaining within a recoverable range.

### Nuances of interpreting breath-hold versus resting-state CVR

The primary dataset for this study utilized breath-hold fMRI to map CVR. Breath holds are a promising approach for mapping CVR that invoke large changes in arterial CO_2_ without the need for external gas delivery, thus reducing equipment costs and increasing participant comfort. During the scan, P_ET_CO_2_ can be measured using a nasal cannula to approximate arterial CO_2_, which allows for CVR calculation in standard units (%BOLD/mmHg). The use of standard units enables CVR comparisons across subjects and scan sessions and facilitates the establishment of normative ranges for healthy CVR values.^12^

We also demonstrated that our main findings were generalizable to resting-state CVR estimates. Resting-state fMRI is a highly feasible method for mapping CVR that exploits natural variations in arterial CO_2_ and may be particularly beneficial for patient populations with difficulties following task instructions. While several studies have validated resting-state CVR estimates,^15,16^ others have suggested that spontaneous breathing fluctuations may not produce sufficient BOLD signal variations for reliable assessment of CVR.^9,66^ Methods for quantitatively mapping CVR (i.e., in units of %BOLD/mmHg) using resting-state data are less common and require P_ET_CO_2_ recordings,^16^ which were not available for our dataset.

We chose to implement the *Rapidtide* toolbox,^59^ which outputs an R^2^ map that reflects the degree to which the blood signal contributes to the BOLD signal in each voxel and is considered a reasonable surrogate for CVR in prior literature examining similar patient groups.^61^ It is important to note that this measure is not identical to conventional CVR since it reflects the shared variance between the probe timeseries and each voxel rather than the magnitude of the voxel’s response. For the purposes of this study and assessing the generalizability of our findings to resting-state data, we considered R² to be a reasonable approximation of CVR. In the future, obtaining P_ET_CO_2_ recordings during resting-state scans could be valuable for refining CVR interpretation without introducing additional patient compliance demands.^16^

### Limitations and future work

One limitation of this study is that we used the same lag range of -12 to +12 seconds for all participants. The lagged general linear model approach used to calculate CVR first accounts for measurement delays and an average vascular transit delay before assessing ±12 second shifts to capture voxel-wise hemodynamic delays. While prior work has suggested lag ranges for stroke populations, their applicability is uncertain due to differences in acquisition and analysis approaches. In healthy participants, lag ranges of ±9 seconds have been successfully used with the lagged general linear model;^18^ we knew we wanted to extend this range slightly to account for potential stroke-related delays. We systematically experimented with different lag ranges and observed that ranges longer than 12 seconds resulted in widespread negative correlations between P_ET_CO_2_ and the BOLD signal (see example in Figure S2). This fixed range may not be appropriate for all participants and brain regions, particularly those near stroke lesions. Future work automating lag-range selection across participants and brain regions will be essential for ensuring accurate CVR interpretation in clinical populations.

Future work should also investigate the physiological basis of negative CVR values observed around vascular lesions in this study. While these negative values may represent a methodological effect of the lag range used, the majority of negative CVR values within vascular lesions were statistically significant (*p*<0.05), suggesting that they may be physiologically meaningful effects (Figure S3). Negative CVR values in peri-infarct regions, evident in the example subjects in Figures 4 and 6, could be attributed to vascular steal, in which blood is diverted from tissue that is already maximally dilated to healthy parenchyma.^67^ Negative CVR values in lesions may arise when cerebral blood volume increases without a corresponding increase in flow,^68^ leading to an accumulation of deoxyhemoglobin within the voxel. In addition, disruption of the blood-brain barrier after stroke can cause vasogenic edema, resulting in increased fluid content within lesion areas.^69^ Negative CVR values in lesions could therefore be caused by increases in blood volume that displace fluid with a higher water density and longer T_2_* than blood. If this effect is sufficiently large, it could overwhelm any positive BOLD effects and lead to a net reduction in BOLD signal.^70,71^ This interpretation is consistent with previous studies investigating negative CVR values in and around brain ventricles, which attributed these effects to a reduction in CSF volume fraction during vasodilation, causing the bright CSF signal to be replaced by a less intensive blood signal.^70–73^

The proposed volume effects driving negative CVR values would be independent of echo time, and thus the persistence of these negative values after ME-ICA denoising in the breath-hold data is surprising. Non-BOLD signal contributions would be anticipated in the SE and ME-OC data, but in the ME-ICA data, we specifically sought to remove ICA components that were not echo-time dependent. This observation may reflect the conservative implementation of ME-ICA used in this study, in which ICA-derived noise components were orthogonalized with respect to signal components, potentially allowing non-BOLD components that are anticorrelated with signal to be retained. We chose to use the conservative ME-ICA approach, rather than using the non-orthogonalized noise components as nuisance regressors, because it has been shown to preserve the BOLD signal variance associated with local CVR responses.^18^ Future studies should specifically evaluate the echo-time dependence of brain regions with negative CVR in order to better understand if these signals arise from BOLD or non-BOLD signal contributions.

In the future, negative CVR could be used to better understand the pathophysiology of stroke and provide insights for predicting functional recovery. In participants with leukoaraiosis, negative CVR values in normal-appearing white matter have been associated with alterations in tissue microstructure and compromised perfusion.^74^ Further investigation is needed to establish whether similar relationships exist in stroke, and the integration of multi-echo fMRI with precision mapping approaches could allow more detailed characterization of negative CVR values. Additionally, considering the timing of the CVR response could help separate flow-related effects related to the local concentration of deoxyhemoglobin from volume-related mechanisms.^71^

Overall, this study demonstrates that multi-echo fMRI is a valuable tool for improving CVR quality in clinical populations. While several important questions remain regarding the interpretation of CVR alterations after stroke and their longitudinal dynamics, multi-echo approaches offer a robust framework for more reliable characterization of these changes.

## Acknowledgements

We are grateful to all the participants for their time and involvement in the study. We thank all the research assistants and students for their help with data collection and manual lesion segmentation. This work was supported by the National Science Foundation (DGE-2234667 to R.G.C.), the UK Medical Research Council (MR/T001402/1), and the National Institute of Neurological Disorders and Stroke (R01NS126509). Infrastructure support was provided by the NIHR Imperial Biomedical Research Centre, the NIHR Imperial Clinical Research Facility, and the Center for Translational Imaging at Northwestern University. The views expressed are those of the authors and not necessarily those of the NSF, NIH, NHS, the NIHR, or the Department of Health and Social Care.

## Author Contributions

Rebecca G. Clements: Conceptualization, Methodology, Software, Formal analysis, Investigation, Data curation, Writing—original draft, Writing—reviewing and editing, Visualization, Project administration. Fatemeh Geranmayeh: Methodology, Resources, Data Curation, Writing—reviewing and editing, Supervision, Funding acquisition. Niamh V. Parkinson: Investigation, Project administration. Miguel Montero: Investigation, Project administration. Sebastian Urday: Resources, Data Curation. Katie Taran: Data curation, Writing—reviewing and editing. Diego A. Caban-Rivera: Data curation, Writing—reviewing and editing. Carson Ingo: Methodology, Resources, Writing—review & editing, Supervision, Project administration, Funding acquisition, Resources. Molly G. Bright: Conceptualization, Methodology, Resources, Writing—review & editing, Supervision, Project administration.

## References

1. Calvo-Imirizaldu M, Solis-Barquero SM, Aramendía-Vidaurreta V, et al. Cerebrovascular Reactivity Mapping in Brain Tumors Based on a Breath-Hold Task Using Arterial Spin Labeling. NMR Biomed 2025; 38: e5317.

2. Silvestrini M, Viticchi G, Falsetti L, et al. The Role of Carotid Atherosclerosis in Alzheimer’s Disease Progression. Journal of Alzheimer’s Disease 2011; 25: 719–726.

3. Van Niftrik CHB, Sebök M, Germans MR, et al. Increased Risk of Recurrent Stroke in Symptomatic Large Vessel Disease with Impaired BOLD Cerebrovascular Reactivity. Stroke 2024; 55: 613–621.

4. Krishnamurthy V, Sprick JD, Krishnamurthy LC, et al. The Utility of Cerebrovascular Reactivity MRI in Brain Rehabilitation: A Mechanistic Perspective. Front Physiol 2021; 12: 642850.

5. Davis TL, Kwong KK, Weisskoff RM, et al. Calibrated functional MRI: Mapping the dynamics of oxidative metabolism. Proc Natl Acad Sci U S A 1998; 95: 1834–1839.

6. Geranmayeh F, Wise RJS, Leech R, et al. Measuring vascular reactivity with breath-holds after stroke: A method to aid interpretation of group-level BOLD signal changes in longitudinal fMRI studies. Hum Brain Mapp 2015; 36: 1755–1771.

7. Nöth U, Meadows GE, Kotajima F, et al. Cerebral vascular response to hypercapnia: Determination with perfusion MRI at 1.5 and 3.0 Tesla using a pulsed arterial spin labeling technique. Journal of Magnetic Resonance Imaging 2006; 24: 1229–1235.

8. Nighoghossian N, Berthezene Y, Meyer R, et al. Assessment of cerebrovascular reactivity by dynamic susceptibility contrast-enhanced MR imaging. J Neurol Sci 1997; 149: 171–176.

9. Pinto J, Bright MG, Bulte DP, et al. Cerebrovascular Reactivity Mapping Without Gas Challenges: A Methodological Guide. Front Physiol 2021; 11: 1711.

10. Lu H, Liu P, Yezhuvath U, et al. MRI Mapping of Cerebrovascular Reactivity via Gas Inhalation Challenges. JoVE (Journal of Visualized Experiments) 2014; e52306.

11. Liu P, Jiang D, Albert M, et al. Multi-vendor and multisite evaluation of cerebrovascular reactivity mapping using hypercapnia challenge. Neuroimage 2021; 245: 118754.

12. Bright MG, Murphy K. Reliable quantification of BOLD fMRI cerebrovascular reactivity despite poor breath-hold performance. Neuroimage 2013; 83: 559–568.

13. Kastrup A, Krüger G, Neumann-Haefelin T, et al. Assessment of cerebrovascular reactivity with functional magnetic resonance imaging: comparison of CO2 and breath holding. Magn Reson Imaging 2001; 19: 13–20.

14. Braban A, Leech R, Murphy K, et al. Cerebrovascular Reactivity Has Negligible Contribution to Hemodynamic Lag After Stroke: Implications for Functional Magnetic Resonance Imaging Studies. Stroke 2023; 54: 1066–1077.

15. Liu P, Liu G, Pinho MC, et al. Cerebrovascular reactivity mapping using resting-state bold functional MRI in healthy adults and patients with moyamoya disease. Radiology 2021; 299: 419–425.

16. Golestani AM, Wei LL, Chen JJ. Quantitative mapping of cerebrovascular reactivity using resting-state BOLD fMRI: Validation in healthy adults. Neuroimage 2016; 138: 147–163.

17. Mandell DM, Han JS, Poublanc J, et al. Quantitative measurement of cerebrovascular reactivity by blood oxygen level-dependent MR imaging in patients with intracranial stenosis: preoperative cerebrovascular reactivity predicts the effect of extracranial-intracranial bypass surgery. AJNR Am J Neuroradiol 2011; 32: 721–727.

18. Moia S, Termenon M, Uruñuela E, et al. ICA-based denoising strategies in breath-hold induced cerebrovascular reactivity mapping with multi echo BOLD fMRI. Neuroimage 2021; 233: 117914.

19. Friston KJ, Williams S, Howard R, et al. Movement-related effects in fMRI time-series. Magn Reson Med 1996; 35: 346–355.

20. Power JD, Mitra A, Laumann TO, et al. Methods to detect, characterize, and remove motion artifact in resting state fMRI. Neuroimage 2014; 84: 320–341.

21. Siegel JS, Power JD, Dubis JW, et al. Statistical improvements in functional magnetic resonance imaging analyses produced by censoring high-motion data points. Hum Brain Mapp 2014; 35: 1981–1996.

22. Caballero-Gaudes C, Reynolds RC. Methods for cleaning the BOLD fMRI signal. Neuroimage 2017; 154: 128–149.

23. Pruim RHR, Mennes M, van Rooij D, et al. ICA-AROMA: A robust ICA-based strategy for removing motion artifacts from fMRI data. Neuroimage 2015; 112: 267–277.

24. Behzadi Y, Restom K, Liau J, et al. A component based noise correction method (CompCor) for BOLD and perfusion based fMRI. Neuroimage 2007; 37: 90–101.

25. Muschelli J, Nebel MB, Caffo BS, et al. Reduction of motion-related artifacts in resting state fMRI using aCompCor. Neuroimage 2014; 96: 22–35.

26. Aranyi C, Opposits G, Nagy M, et al. Population-Level Correction of Systematic Motion Artifacts in fMRI in Patients with Ischemic Stroke. Journal of Neuroimaging 2017; 27: 397–408.

27. Patriat R, Reynolds RC, Birn RM. An improved model of motion-related signal changes in fMRI. Neuroimage 2017; 144: 74–82.

28. Poser BA, Versluis MJ, Hoogduin JM, et al. BOLD contrast sensitivity enhancement and artifact reduction with multiecho EPI: Parallel-acquired inhomogeneity-desensitized fMRI. Magn Reson Med 2006; 55: 1227–1235.

29. DuPre E, Salo T, Ahmed Z, et al. TE-dependent analysis of multi-echo fMRI with *tedana*. J Open Source Softw 2021; 6: 3669.

30. Kundu P, Inati SJ, Evans JW, et al. Differentiating BOLD and non-BOLD signals in fMRI time series using multi-echo EPI. Neuroimage 2012; 60: 1759–1770.

31. Dipasquale O, Sethi A, Lagan MM, et al. Comparing resting state fMRI de-noising approaches using multi- and single-echo acquisitions. PLoS One 2017; 12: e0173289.

32. Reddy NA, Zvolanek KM, Moia S, et al. Denoising task-correlated head motion from motor-task fMRI data with multi-echo ICA. Imaging Neuroscience 2024; 2: 1–30.

33. Fernandez B, Leuchs L, Sämann PG, et al. Multi-echo EPI of human fear conditioning reveals improved BOLD detection in ventromedial prefrontal cortex. Neuroimage 2017; 156: 65–77.

34. Cohen AD, Wang Y. Improving the Assessment of Breath-Holding Induced Cerebral Vascular Reactivity Using a Multiband Multi-echo ASL/BOLD Sequence. Scientific Reports 2019 9:1 2019; 9: 5079-.

35. Lynch CJ, Power JD, Scult MA, et al. Rapid Precision Functional Mapping of Individuals Using Multi-Echo fMRI. Cell Rep 2020; 33: 108540.

36. Seto E, Sela G, McIlroy WE, et al. Quantifying Head Motion Associated with Motor Tasks Used in fMRI. Neuroimage 2001; 14: 284–297.

37. Gruia DC, Trender W, Hellyer P, et al. IC3 protocol: a longitudinal observational study of cognition after stroke using novel digital health technology. BMJ Open 2023; 13: e076653.

38. Friston KJ, Fletcher P, Josephs O, et al. Event-Related fMRI: Characterizing Differential Responses. Neuroimage 1998; 7: 30–40.

39. Zvolanek KM, Moia S, Dean JN, et al. Comparing end-tidal CO2, respiration volume per time (RVT), and average gray matter signal for mapping cerebrovascular reactivity amplitude and delay with breath-hold task BOLD fMRI. Neuroimage 2023; 272: 120038.

40. Clements RG, Zvolanek KM, Reddy NA, et al. Quantitative mapping of cerebrovascular reactivity amplitude and delay with breath-hold BOLD fMRI when end-tidal CO2 quality is low. Imaging Neuroscience 2025; 3: 2025.

41. Jenkinson M, Beckmann CF, Behrens TEJ, et al. FSL. Neuroimage 2012; 62: 782–790.

42. Cox RW. AFNI: Software for Analysis and Visualization of Functional Magnetic Resonance Neuroimages. Computers and Biomedical Research 1996; 29: 162–173.

43. Cox RW, Hyde JS. Software Tools for Analysis and Visualization of fMRI Data. Epub ahead of print 1997. DOI: 10.1002/(SICI)1099-1492(199706/08)10:4/5.

44. Zhang Y, Brady M, Smith S. Segmentation of brain MR images through a hidden Markov random field model and the expectation-maximization algorithm. IEEE Trans Med Imaging 2001; 20: 45–57.

45. Jenkinson M, Smith S. A global optimisation method for robust affine registration of brain images. Med Image Anal 2001; 5: 143–156.

46. Jenkinson M, Bannister P, Brady M, et al. Improved Optimization for the Robust and Accurate Linear Registration and Motion Correction of Brain Images. Neuroimage 2002; 17: 825–841.

47. Jenkinson M. Improving the registration of B0- disorted EPI images using calculated cost function weights. In: Conf. on Functional Mapping of the Human Brain. 2004.

48. Jezzard P. Correction of geometric distortion in fMRI data. Neuroimage 2012; 62: 648651.

49. Ahmed Z, Bandettini PA, Bottenhorn KL, et al. ME-ICA/tedana: 25.1.1a2. Epub ahead of print 15 December 2025. DOI: 10.5281/ZENODO.17943951.

50. Kundu P, Brenowitz ND, Voon V, et al. Integrated strategy for improving functional connectivity mapping using multiecho fMRI. Proc Natl Acad Sci U S A 2013; 110: 16187–16192.

51. tedana: TE Dependent ANAlysis — tedana 25.1.0 documentation, https://tedana.readthedocs.io/en/stable/# (2025, accessed 19 November 2025).

52. Reddy NA, Medina MC, Northrop JN, et al. Impact of multi-echo ICA modeling decisions on motor-task fMRI analysis. bioRxiv 2025; 2025.11.25.690440.

53. Moia S, Vigotsky AD, Zvolanek KM. phys2cvr: A tool to compute Cerebrovascular Reactivity maps and associated lag maps. Epub ahead of print 1 June 2024. DOI: 10.5281/ZENODO.7336002.

54. Moia S, Stickland RC, Ayyagari A, et al. Voxelwise optimization of hemodynamic lags to improve regional CVR estimates in breath-hold fMRI. In: Proceedings of the Annual International Conference of the IEEE Engineering in Medicine and Biology Society, EMBS. Institute of Electrical and Electronics Engineers Inc., 2020, pp. 1489–1492.

55. Power JD, Barnes KA, Snyder AZ, et al. Spurious but systematic correlations in functional connectivity MRI networks arise from subject motion. Neuroimage 2012; 59: 2142–2154.

56. Smyser CD, Inder TE, Shimony JS, et al. Longitudinal Analysis of Neural Network Development in Preterm Infants. Cerebral Cortex 2010; 20: 2852–2862.

57. Schmal C, Myung J, Herzel H, et al. Moran’s I quantifies spatio-temporal pattern formation in neural imaging data. Bioinformatics 2017; 33: 3072–3079.

58. Rey SJ, Anselin L. PySAL: A Python Library of Spatial Analytical Methods. Handbook of Applied Spatial Analysis 2010; 175–193.

59. Frederick B. rapidtide. Epub ahead of print 2024. DOI: 10.5281/zenodo.814990.

60. Gong J, Stickland RC, Bright MG. Hemodynamic timing in resting-state and breathing-task BOLD fMRI. Neuroimage 2023; 274: 120120.

61. Donahue MJ, Strother MK, Lindsey KP, et al. Time delay processing of hypercapnic fMRI allows quantitative parameterization of cerebrovascular reactivity and blood flow delays. Journal of Cerebral Blood Flow and Metabolism 2016; 36: 1767–1779.

62. Renger E, Hauser TK, Klose U, et al. Assessing and Improving the Reproducibility of Cerebrovascular Reactivity Evaluations in Healthy Subjects Using Short-Breath-Hold fMRI. Diagnostics 2025, Vol 15, Page 1946 2025; 15: 1946.

63. Elliott ML, Knodt AR, Hariri AR. Striving toward translation: strategies for reliable fMRI measurement. Trends Cogn Sci 2021; 25: 776–787.

64. Stickland RC, Zvolanek KM, Moia S, et al. A practical modification to a resting state fMRI protocol for improved characterization of cerebrovascular function. Neuroimage 2021; 239: 118306.

65. Lipp I, Murphy K, Caseras X, et al. Agreement and repeatability of vascular reactivity estimates based on a breath-hold task and a resting state scan. Neuroimage 2015; 113: 387.

66. De Vis JB, Bhogal AA, Hendrikse J, et al. Effect sizes of BOLD CVR, resting-state signal fluctuations and time delay measures for the assessment of hemodynamic impairment in carotid occlusion patients. Neuroimage 2018; 179: 530–539.

67. Poublanc J, Han JS, Mandell DM, et al. Vascular Steal Explains Early Paradoxical Blood Oxygen Level-Dependent Cerebrovascular Response in Brain Regions with Delayed Arterial Transit Times. Cerebrovasc Dis Extra 2013; 3: 55.

68. Siemund R, Cronqvist M, Andsberg G, et al. Cerebral Perfusion Imaging in Hemodynamic Stroke: Be Aware of the Pattern. Interventional Neuroradiology 2009; 15: 385.

69. Kim BJ, Kang HG, Kim H-J, et al. Magnetic Resonance Imaging in Acute Ischemic Stroke Treatment. J Stroke 2014; 16: 131.

70. Thomas BP, Liu P, Aslan S, et al. Physiologic underpinnings of negative BOLD cerebrovascular reactivity in brain ventricles. Neuroimage 2013; 83: 505–512.

71. Bright MG, Bianciardi M, de Zwart JA, et al. Early anti-correlated BOLD signal changes of physiologic origin. Neuroimage 2014; 87: 287–296.

72. Blockley NP, Driver ID, Francis ST, et al. An improved method for acquiring cerebrovascular reactivity maps. Magn Reson Med 2011; 65: 1278–1286.

73. Bianciardi M, Fukunaga M, Van Gelderen P, et al. Negative BOLD-fMRI signals in large cerebral veins. Journal of Cerebral Blood Flow & Metabolism 2010; 31: 401.

74. Sam K, Peltenburg B, Conklin J, et al. Cerebrovascular reactivity and white matter integrity. Neurology 2016; 87: 2333.

