## Supplementary Material for "Multi-echo BOLD fMRI improves cerebrovascular reactivity estimates in stroke"

### Methods

##### *Identification of acute infarcts in resting-state dataset*

To create infarct masks for each participant in the resting-state dataset, we used their clinical scans acquired at acute stroke onset at Northwestern Memorial Hospital on a 3T Siemens Skyra MRI scanner. Infarct diagnosis was confirmed with the following diffusion-weighted MRI (dMRI) protocol: TR = 8200 ms, TE = 89 ms, 50 axial slices, FOV = 240 x 240 $mm^2$, voxel resolution = 1.25x1.25x3 $mm^3$. A trained neuroradiologist identified each infarct via the diffusion weighted image (DWI) and the apparent diffusion coefficient (ADC) maps in which diffusivities were determined to be less than 620 x $10^{-6}$ $mm^2/s$.[1]

##### *Pre-processing of resting-state dataset*

The T1-weighted scans were brain extracted and used to generate gray and white matter masks (partial volume threshold = 0.5). The infarct masks identified using the acute DWI were transformed to T1-space and subtracted from the gray and white matter masks, and the resulting normal-appearing gray and white matter masks were transformed to functional space (*flirt*, FSL). Head motion realignment (*3dVolReg*, AFNI) was performed with the SBRef as the reference volume and then distortion correction was performed using *topup* (FSL). Next, a brain mask was generated using the distortion-corrected SBRef and used to brain extract each echo. As done for the breath-hold dataset, the pre-processed second echo was used as a surrogate for SE data. The ME-OC dataset was generated using *tedana*[2,3], and ME-ICA was performed using the *tedana* decision tree described for the breath-hold dataset.

##### *Denoising of resting-state dataset*

Denoising was performed using AFNI's 3dREMLfit. For the SE and ME-OC approaches, we removed the 6 demeaned motion parameters obtained during volume registration and their temporal derivatives and Legendre polynomials up to 4$^{th}$ order. For the ME-ICA approach, we additionally removed the rejected ICA components orthogonalized to the mean time series in gray matter in the non-lesioned hemisphere, accepted ICA components, motion parameters and their derivatives, and Legendre polynomials. Each denoised dataset was smoothed using a 4mm FWHM Gaussian kernel.

##### *Visualization of 3 representative resting-state subjects*

For the visualization of 3 representative resting-state subjects in this paper, we registered their $R^2$ and r maps to MNI space using the FSL 1mm MNI template resampled to 2.5 mm resolution (FLIRT and FNIRT, FSL). To visualize white matter injuries for these subjects, in addition to the manually drawn infarct mask, hyperintensities were segmented from the FLAIR image using the lesion prediction algorithm (LPA) as implemented in the Lesion Segmentation Toolbox (LST; version 3.0.0; https://www.statistical-modelling.de/lst.html) for SPM.[4,5] The T1-weighted image was used as the anatomical reference for co-registration within SPM prior to lesion segmentation. The resulting LPA-derived lesion probability map was converted to a binary hyperintensity mask, registered to the MNI template, and overlaid on each example CVR map alongside the manually drawn infarct mask.

#### Results

##### *Sample Size Justification*

After excluding participants with low-quality $P_{ET}CO_2$ timeseries, the breath-hold dataset consisted of 36 participants. We assessed the SE, ME-OC, and ME-ICA methods using both quantitative metrics and qualitative observations, and it is challenging to reduce this study to a single primary outcome for a power analysis. However, one primary outcome of this study is the percentage of significant gray matter voxels in gray matter, comparing SE to ME-ICA. The effect size for this measure was 0.79, and with $\alpha = 0.05$ and 36 participants, the resulting power for a two-tailed test was 0.996, indicating that our sample size is sufficient to detect differences.

#### Discussion

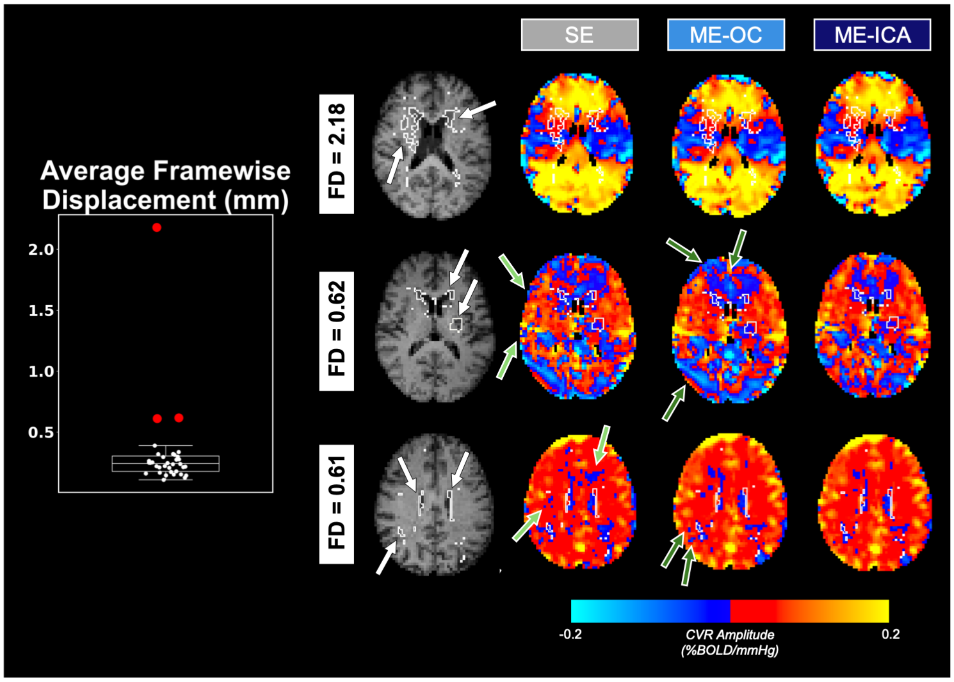


**Figure S1.** Average framewise displacement values (left) and CVR maps generated using SE, ME-OC, and ME-ICA for the 3 participants with the highest average framewise displacement values. Lesions and white matter hyperintensities are outlined in white and indicated by white arrows on the T1-weighted anatomical scans. Light green arrows (second column of brain maps) indicate non-lesion and non-white matter hyperintensity areas with fewer negative CVR values using ME-OC compared to SE. Dark green arrows (third column of brain maps) indicate non-lesion and non-white matter hyperintensity areas with fewer negative CVR values using ME-ICA compared to ME-OC.

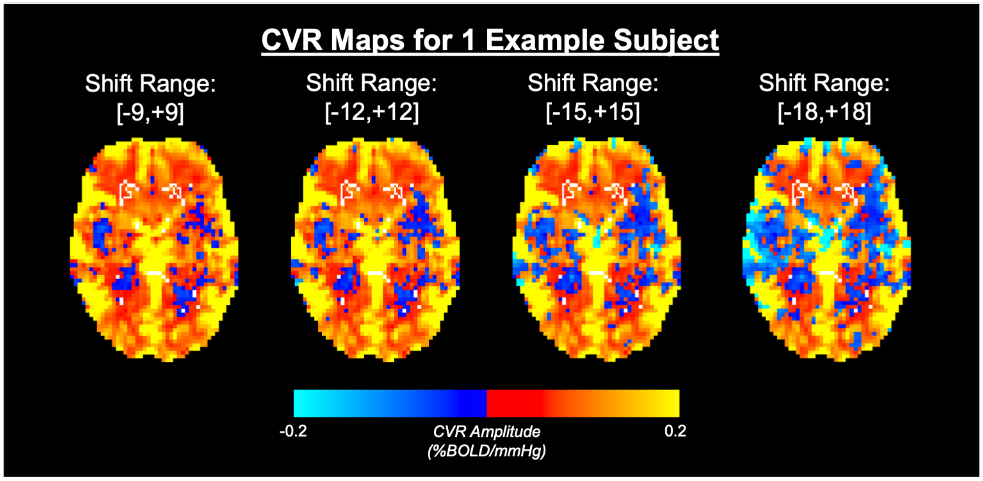


**Figure S2**. Lag maps generated using varying shift ranges for 1 example subject. The maps generated using the [-15,+15] and [-18,+18] shift ranges have widespread negative CVR values in normal-appearing gray and white matter that are not seen in the maps generated using the [-9,+9] and [-12,+12] shift ranges. Vascular lesions are outlined in white.

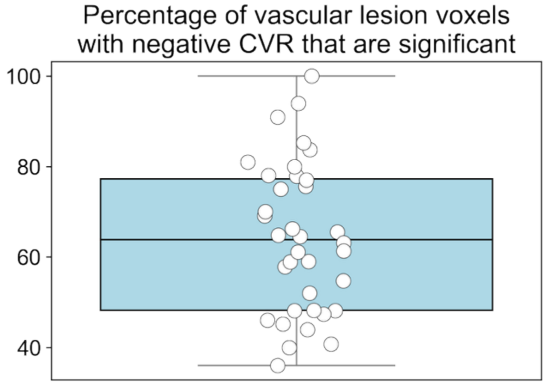


**Figure S3**. Percentage of vascular lesion voxels with negative CVR that are statistically significant. Significant voxels were identified as having absolute T-statistics greater than 1.96 (corresponding to $p<0.05$). Each dot represents a subject. This plot shows that, on average, 64% of negative CVR voxels within vascular lesions were statistically significant.

#### References

1. Purushotham A, Campbell BCV, Straka M, et al. Apparent diffusion coefficient threshold for delineation of ischemic core. *International Journal of Stroke* 2015; 10: 348–353.

2. Ahmed Z, Bandettini PA, Bottenhorn KL, et al. ME-ICA/tedana: 25.1.0. *Zenodo*. Epub ahead of print 25AD. DOI: 10.5281/ZENODO.17379477.

3. DuPre E, Salo T, Ahmed Z, et al. TE-dependent analysis of multi-echo fMRI with *tedana*. *J Open Source Softw* 2021; 6: 3669.

4. Schmidt P. Bayesian inference for structured additive regression models for large-scale problems with applications to medical imaging. *MU München: Faculty of Mathematics, Computer Science and Statistics*. Epub ahead of print 19 January 2017. DOI: 10.5282/EDOC.20373.

5. Schmidt P, Gaser C, Arsic M, et al. An automated tool for detection of FLAIR-hyperintense white-matter lesions in Multiple Sclerosis. *Neuroimage* 2012; 59: 3774–3783.
